# Visual recency bias is explained by a mixture model of internal representations

**DOI:** 10.1101/228973

**Authors:** Kristjan Kalm, Dennis Norris

## Abstract

Human bias towards more recent events is a common and well-studied phenomenon. Recent studies in visual perception have shown that this recency bias persists even when past events contain no information about the future. Reasons for this suboptimal behaviour are not well understood and the internal model that leads people to exhibit recency bias is unknown. Here we use a well-known orientation estimation task to frame the human recency bias in terms of incremental Bayesian inference. We show that the only Bayesian model capable of explaining the recency bias relies on a weighted mixture of past states. Furthermore, we suggest that this mixture model is a consequence of participants’ failure to infer a model for data in visual short term memory, and reflects the nature of the internal representations used in the task.

## Introduction

In a rapidly changing world our model of the environment needs to be continuously updated. Often recent information is a better predictor of the environment than the more distant past (Anderson & Milson, 1989; Anderson & Schooler, 1991): for example, the location of a moving object is better predicted by its location one second ago than a minute ago. However, human observers seem to rely on recent experience even when it provides no information about the future at all (Fischer & Whitney, 2014; Cicchini, Anobile, & Burr, 2014; Burr & Cicchini, 2014; Fritsche, Mostert, & de Lange, 2017; Liberman, Fischer, & Whitney, 2014). Such recency bias seems to be domain-general and not constrained to a particular task or feature dimension (Kiyonaga, Scimeca, Bliss, & Whitney, 2017). Why should this be so, and what can it tell us about the mechanisms of perception and memory?

The most extensive quantitative data on the human recency bias comes from a study of visual orientation estimation by Fischer and Whitney (2014). In that study participants were presented with a randomly oriented grating (Gabor) on each trial and asked to report the orientation by adjusting a bar using the arrow keys (Fig 1A).

**Figure 1:**
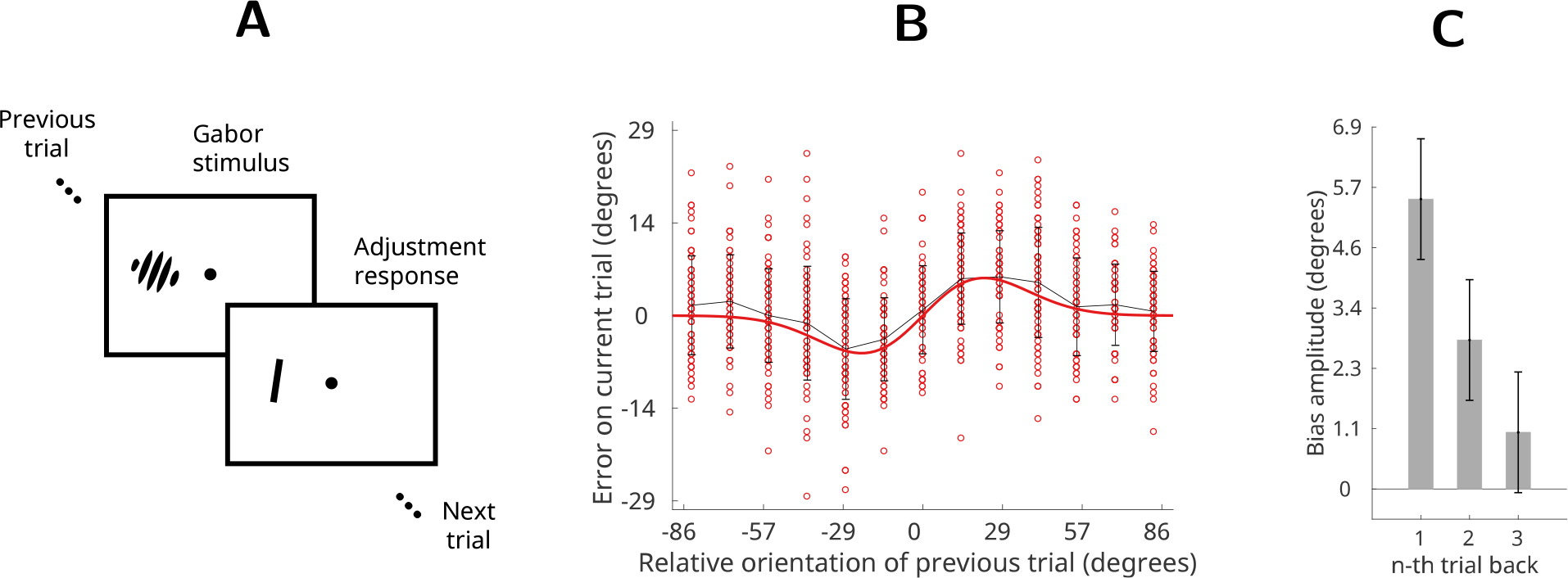
Orientation estimation task (Fischer & Whitney, 2014) (A) Participants observed randomly oriented Gabor stimuli and reported the orientation of each Gabor by adjusting a response bar. Stimuli were presented for 500 ms and separated in time by 5s. (B) Single subject’s errors (red dots) as a function of the relative orientation of the previous trial. Gray line is average error; red line shows a first derivative of a Gaussian (DoG) curve fit to the data. (C) Average recency bias amplitude across participants computed for stimuli presented one, two and three trials back from the present. Error bars represent ±1 standard deviation of the bootstrapped distribution.

Participants’ error distributions revealed that although responses were centred on the correct orientations over the course of the entire experiment, on a trial-by-trial basis the reported orientation was systematically (and precisely) biased in the direction of the orientation seen on the previous trial. For example, when the Gabor on the previous trial was oriented more clockwise than the Gabor on the present trial, participants perceived the present Gabor as being tilted more clockwise than its true orientation (Fig 1B).

Since the orientations of the stimuli were generated randomly in this task the recency bias indicates that participants are not behaving optimally. In other words, the previous trial contained no information about the next trial and hence the optimal model would consider all orientations as equally likely in the future. In this case the participants’ error distributions would simply be proportional to the sensory noise and always centred around the true stimulus value (Fig 2A, top row). However, here participants assume a model of the environment where past states are informative about the future (Fig 2A, bottom row), which is clearly false.

In the current study we use the orientation estimation task (Fischer & Whitney, 2014) to investigate what is the participants’ model of the environment that gives rise to the recency bias. We frame this question in terms of sequential Bayesian inference which allows us to test hypotheses about the participant’s model of the environment at any trial given sensory information (orientation of the Gabor) and the recorded response (Fig 2, see also *Bayesian orientation estimation* in *Supporting Information*). We test three alternative hypotheses about the model behind the recency bias which are all formulated as sequential Bayesian inference models so that they can be directly compared to each other.

**Figure 2:**
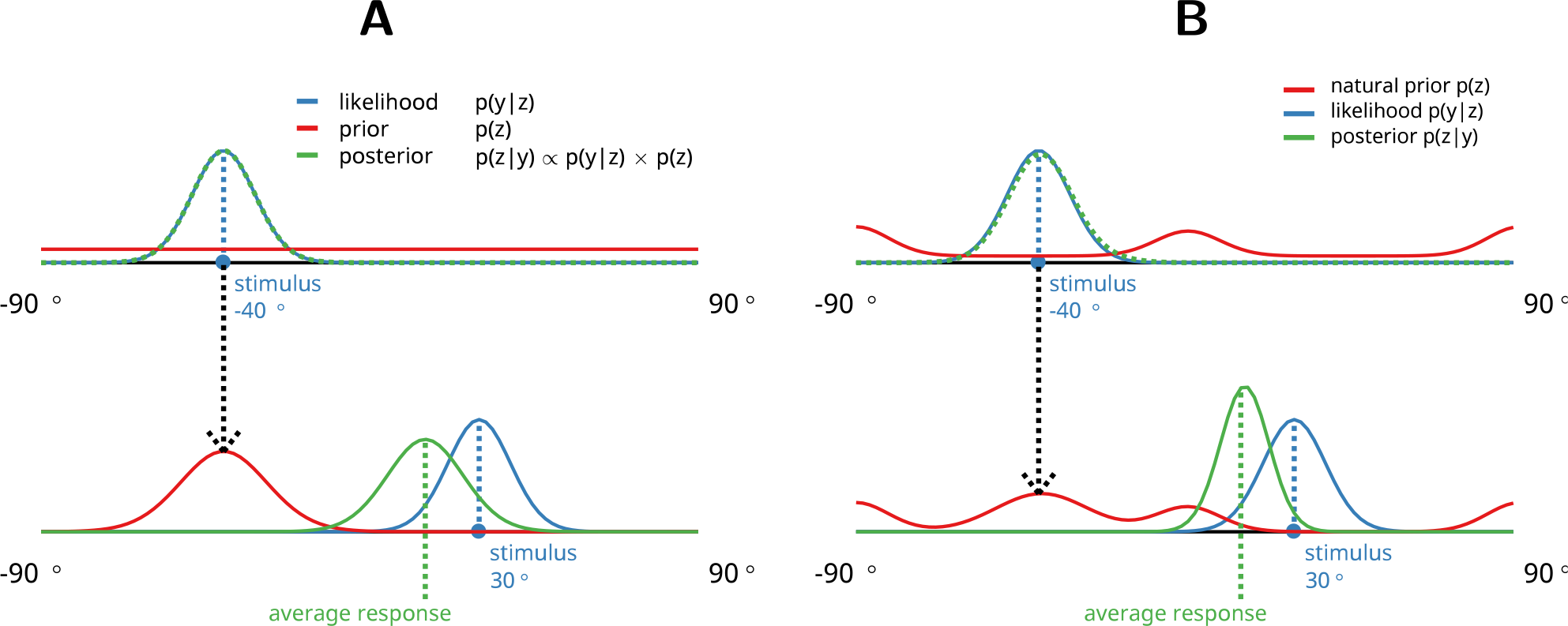
Bayesian orientation estimation and recency bias. Participant’s estimate of the orientation (*p*(*z|y*), green line) combines sensory evidence (*p*(*y|z*), blue line) with prior expectation (*p*(*z*), red line). Participant’s response can be thought of as a sample from the posterior distribution. All distributions are Von Mises since orientation is a circular variable. (A) Top row: for optimal behaviour participant’s prior should be flat and posterior equal to the sensory evidence. Bottom row: Recency bias occurs when information about previous stimuli (orientation estimate at trial *n* − 1, green line in top row) is transferred to the prior expectation about the next stimulus (red line, bottom row). Here the prior for trial *t* is just the posterior from previous trial *t* − 1. (B) Natural prior model. Top row: participants prior is based on the statistics of the natural environment (Girshick et al., 2011). Bottom row: participants prior is a mixture of the previous stimulus and the natural statistics.

### Von Mises filter

First, we test the hypothesis that participants assume that the current state of the environment is the best guess about it’s future. This *identity model* is the simplest Bayesian incremental updating model (a Bayesian filter) that can plausibly represent the orientation estimation task. Bayesian filters (such as the Kalman filter, Kalman & Bucy, 1961) are widely used in explaining human behaviour and have been previously proposed to explain the temporal continuity effects in perception (Burr & Cicchini, 2014; Wolpert & Ghahramani, 2000; Rao, 1999).

Here we use the circular approximation of the Kalman filter called the Von Mises filter (VMF) where the latent state and measurement noise are distributed according to Von Mises and not Gaussian distributions (Kurz, Gilitschenski, & Hanebeck, 2016; Marković & Petrović, 2009). An example of a simple VMF is depicted on Fig 2A, where the prediction *p*(*z_t_*) at the bottom row is derived from the previously estimated posterior distribution 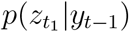, or in other words, the latent state transition model is identity. See *Von Mises filter* in *Methods* for details.

### Natural prior model

A simple identity model as outlined above ignores the fact that people’s orientation judgements are more accurate at cardinal orientations, reflecting the statistics of contours in natural environment (Girshick et al., 2011). Such bias suggests that the observer’s internal model matches the environment, a hallmark of Bayesian optimality. Fig 3 depicts orientation statistics of a natural image and participants’ orientation sensitivity extracted from a behavioural task (Girshick et al., 2011). Hence we can supplement the identity model with a *natural prior* so that participants modulate the identity prediction by taking into account the natural statistics of orientations in the environment.

Since the size of the recency bias in the task was independent of stimulus orientation (Fischer & Whitney, 2014) we can rule out a static natural prior in advance. Instead, we assume here that the prior is a mixture of the stimulus on the previous trial and the natural prior distribution. An illustration of a single step in the natural prior model is depicted on Fig 2B, where the prediction *z*_*t*_ is equal to the mixture of the previous stimulus and a static natural prior. See *Natural prior model* in *Methods* for details.

### Mixture model

Last, we test the hypothesis that participants’ predictions incorporate information from multiple past trials. Such a *mixture model* assumes that the participant’s model of the environment is a mixture of multiple past states so that more recent states contribute more than older ones. The two previous hypotheses both assume that the model of the environment *p*(*z*) is inferred only based on the previous latent state. Contrastingly, the human recency bias clearly extends beyond the previous state - it is greatest for the most recent state and decays for each further state into the past (Fig 1C). In order to model such time-decaying recency bias over several past states we modify the VMF so that its prior distribution reflects a time-decaying mixture of information from multiple previous trials. Fig 4 illustrates the evolution of the latent state *p*(*z*) in a mixture model over 4 trials. Importantly, such mixture distribution is computed by a fixed sampling step (Kalm, 2017) which results in a computationally first-order Markovian model which has the same number of parameters and model complexity as the natural prior model described above. See *Mixture model* in *Methods* for details.

**Figure 3:**
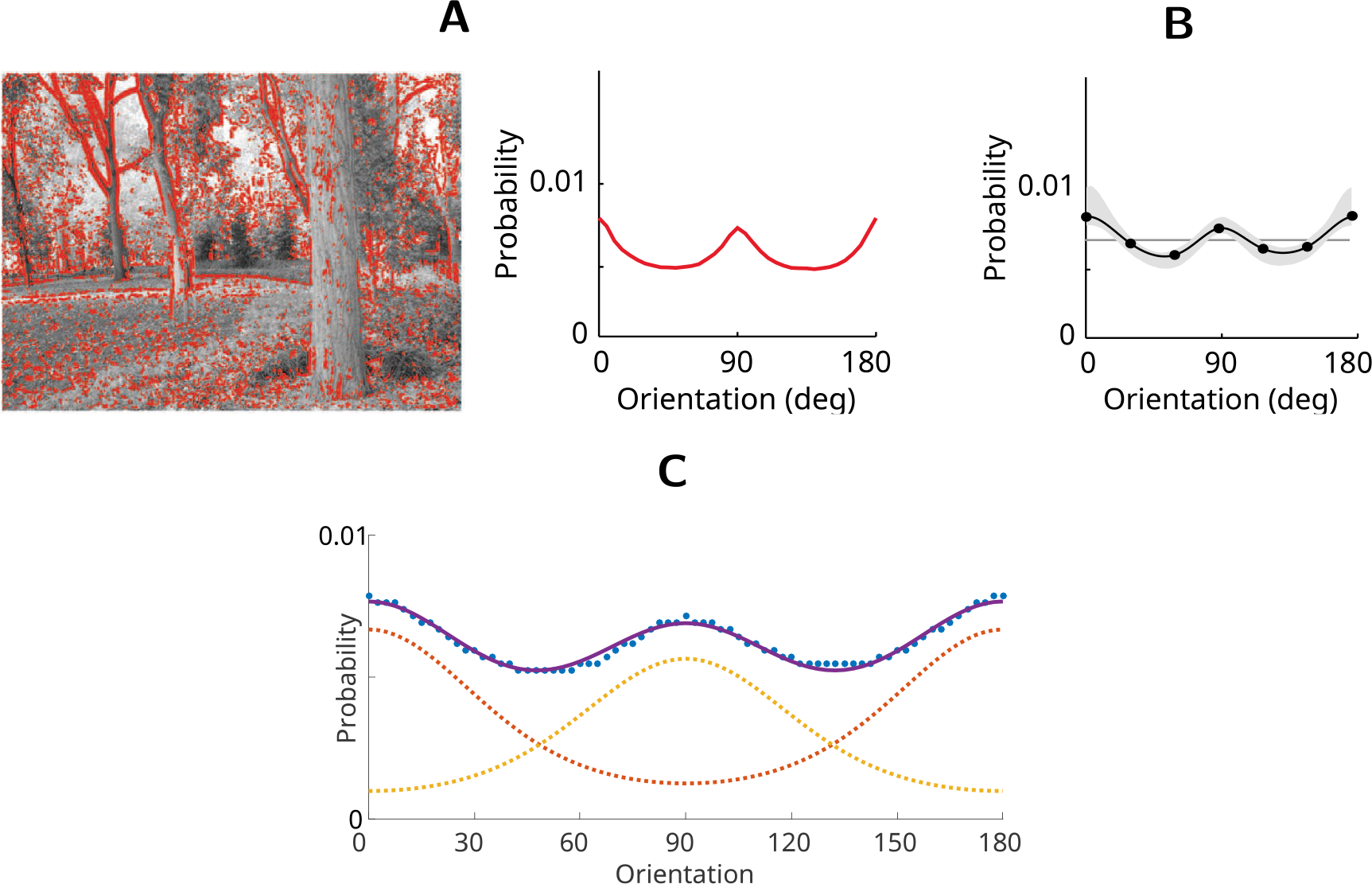
Natural prior for orientation (Girshick et al., 2011). (A) A natural image (left) and a distribution of contour orientations extracted from the image. (B) Average prior distribution of orientations across all participants estimated with a noisy orientation judgement task. The grey error region shows ±1 standard deviation of 1,000 bootstrapped estimated priors. (C) Observers’ average prior as reported by Girshick et al., 2011 (dotted blue line) represented as a mixture of two Von Mises distributions (solid blue line), which has two components peaking at cardinal orientations (dotted yellow and red lines).

Note that we can a priori rule out approaches which track the average orientation or some other summary statistic (Hubert-Wallander & Boynton, 2015; Dubé, Zhou, Kahana, & Sekuler, 2014) since with random stimuli they would all be uninformative about the past (however, see Manassi, Liberman, Chaney, & Whitney, 2017 for sequential dependencies in summary statistical judgments themselves).

**Figure 4:**
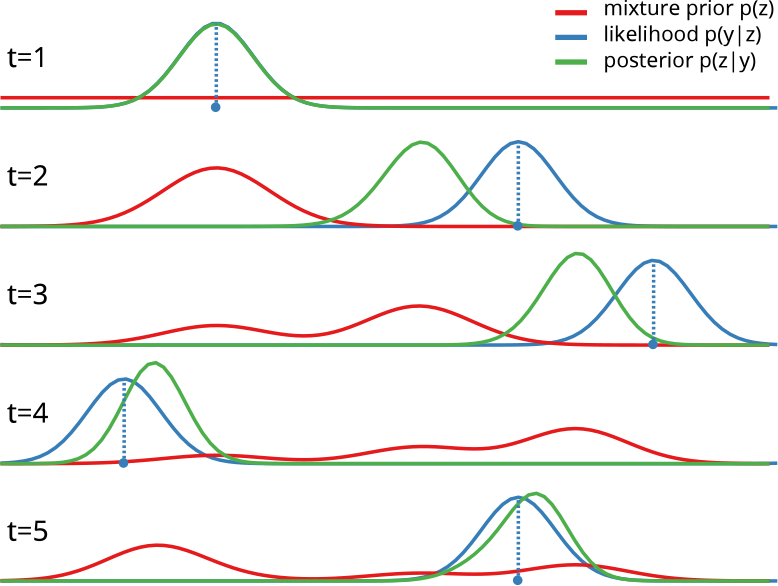
Mixture model. Evolution of the mixture latent state over five trials.

## Methods

### Von Mises filter

Here we use the circular approximation of the Kalman filter called the Von Mises filter (VMF) where the latent state and measurement noise are distributed according to Von Mises and not Gaussian distributions,

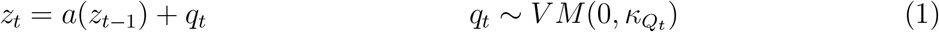

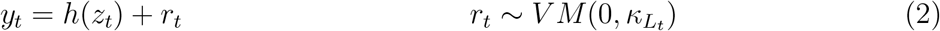

where *κ_Q_* and *κ_L_* are latent state and measurement noise concentration terms respectively. An example of a simple VMF is depicted on Fig 2B, where both state transition and measurement models are identity functions and state noise is zero, resulting in a model where the predicted state *z_t_* is equal to the previously estimated posterior distribution 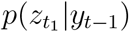. The posterior distribution, being a product of two Von Mises distributions and representing participant’s estimate, therefore also approximates Von Mises (see *Product of two von Mises distributions* in *Supporting Information* for details):

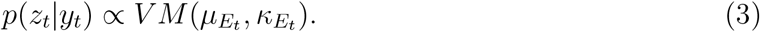

This allows us to define recency bias on any trial *t* as the distance which posterior mean 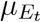 has moved away from the presented stimulus *y_t_* towards some previous stimulus value *y_t-n_*. Such estimation error represents the systematic shift in participants’ responses since the internal estimate of the perceived orientation (Eq 3) is not centred around the presented stimulus *y_t_* (Fig 2). The value of the estimation error, as a distance between the posterior mean and stimulus value, can be easily derived from the properties of Von Mises product:

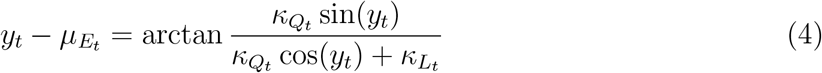

Importantly, this estimation error function (Eq 4) allows us to describe the possible space of recency biases by mapping the systematic shift of the estimation error towards previously observed orientations (Fig 5).

**Figure 5:**
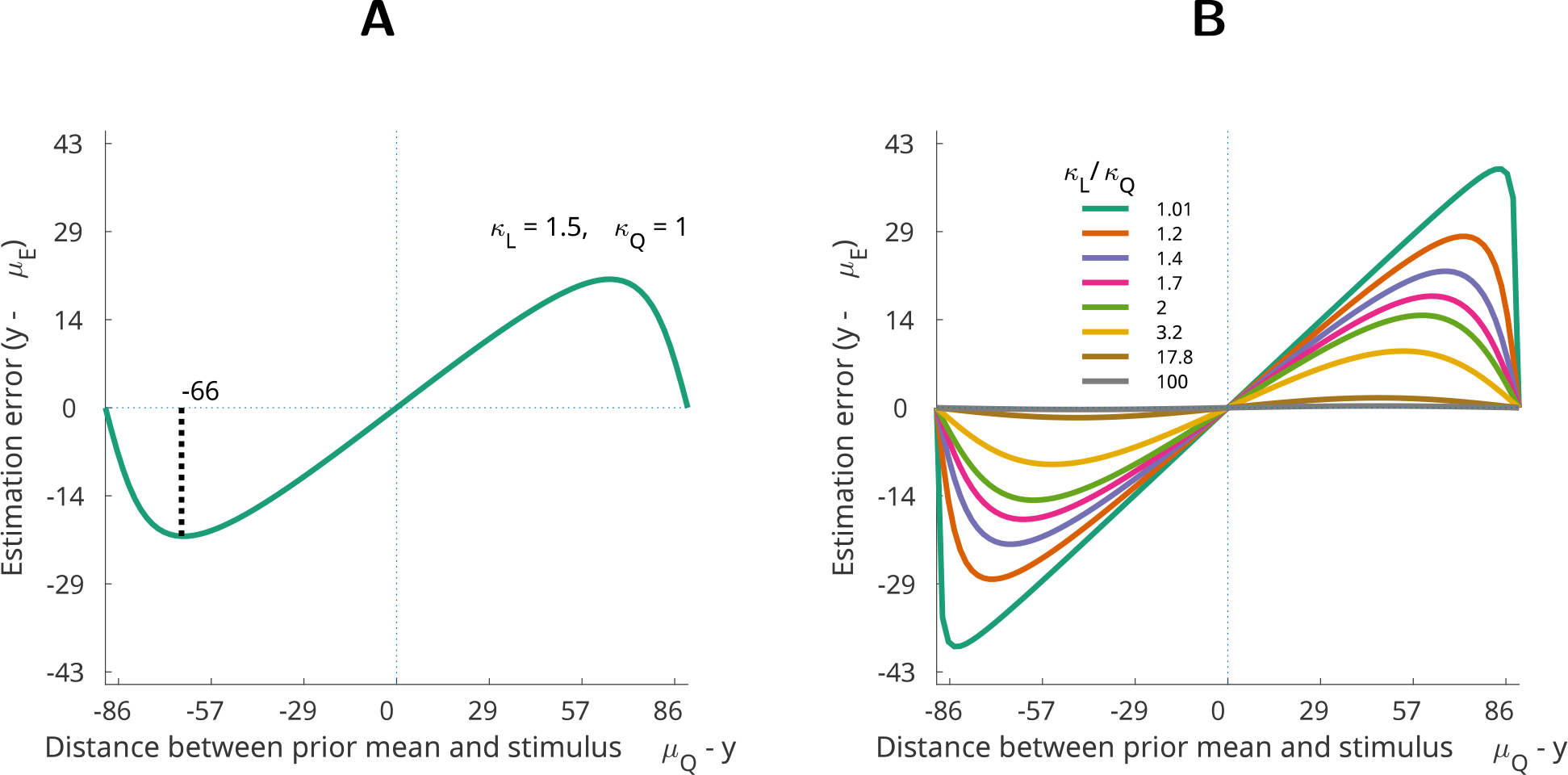
Recency bias with Von Mises distributions. (A) Example of Von Mises recency bias with fixed prior and likelihood parameters (*κ_L_* = 1.5, *κ_Q_* = 1, Eq 4). Recency bias is greatest when the distance between the prior mean and stimulus (x-axis) is ca 66 degrees (1.15 radians) (black dotted line). (B) The shape of the recency bias depends on the ratio of likelihood to prior concentration 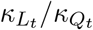 (differently coloured lines). See *Von Mises filter properties* in *Supporting Information* for details.

Such mapping of all possible shapes of the recency bias (Fig 5B) reveals that when Von Mises distributions are used for Bayesian inference the recency bias always peaks more than halfway through the x-axis (Fig 5, for a proof, see *Von Mises filter properties* in *Supporting Information).* This property means that the VMF cannot even theoretically yield a DoG-like recency bias shape as observed with human participants (Fig 1B).

#### Model parameters

To model the perceptual noise around the stimulus value (Eq 2) we used a fixed concentration parameter for the likelihood function (*κ_L_*), which was chosen so as to produce the just noticeable difference (JND) values matching human data from Fischer and Whitney (2014) (average JND was 5.39°, hence *σ* = 3.8113° and *κ_L_* = 0.0688). The concentration parameter for the state noise *κ_Q_* was a free parameter optimised to minimise the distance between the simulated data and the average observed subjects’ response (see *Model fitting and parameter optimisation* in *Supporting Information*).

### Natural prior model

We modified the VMF so that instead of predicting the next state based on the current one (identity model) we assume that everything else being equal, cardinal orientations are more likely than oblique ones and reflect this in our prediction. We can do this by using the natural prior distribution function as the state transition model which changes the predictive prior distribution *p*(*z_t_*|*z*_*t*-1_) on trial *t* from unimodal Von Mises to bimodal non-parametric distribution. For this purpose we model the average observers’ prior as reported by Girshick et al. (2011) as a mixture of two Von Mises probability distributions, which has two components peaking at cardinal orientations (solid blue line on Fig 3C):

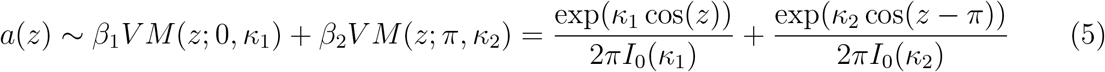

We can now specify the equations for the Bayesian filter (Eq 1,2) with the natural prior with a state transition model *a*(*z*_*t*−1_) is a bimodal mixture peaking at cardinal orientations (Eq 5, Fig 3C), the measurement model is identity, and the noise for both is additive Von Mises.

However, if the prior would always predict cardinals over obliques, we would only observe recency bias for the trials which were preceded by orientations close to cardinal angles. Since the size of the recency bias in the behavioural experiment was independent of stimulus orientation (Fischer & Whitney, 2014) we can rule out a fixed natural prior (Eq 5) in advance. Instead, we assume here that the prior is a mixture of the natural prior and the previous posterior.

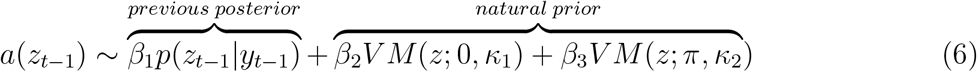

This leads to a prediction which is still biased towards the previous trial but mixed with the natural prior (Fig 6). In general terms, we assume here that participants have both a bias towards previous orientations and natural statistics of the environment (Fig 3). Importantly, such multi-modal prior means that participants estimates (posterior distribution) are also not Von Mises which allows for recency bias curves qualitatively different from Von Mises ones.

**Figure 6:**
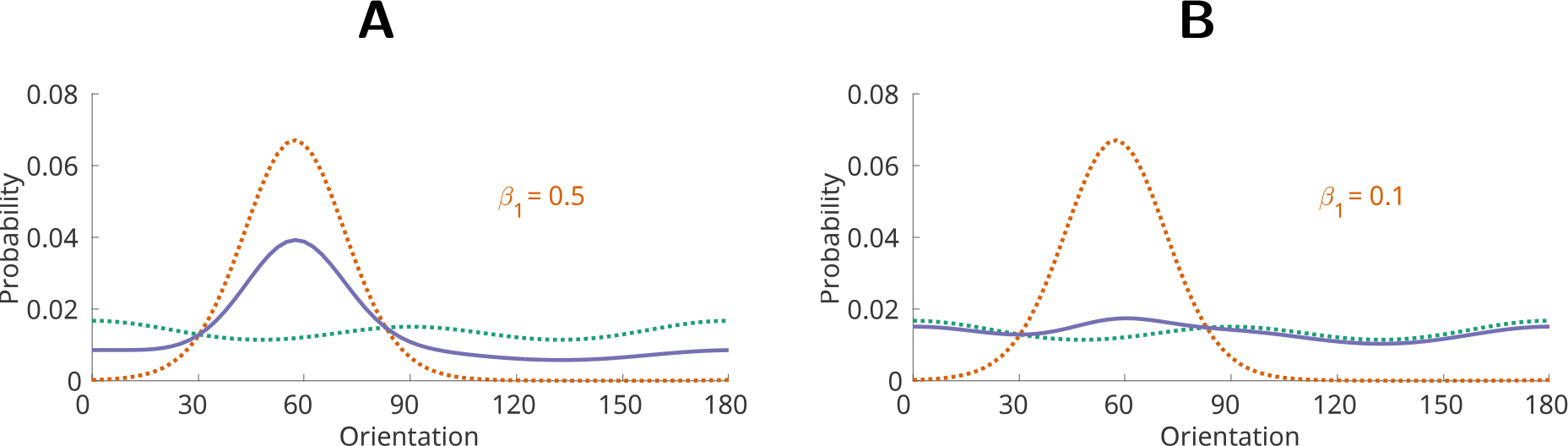
Prior distribution as a mixture between the previous posterior *p*(*z*_*t*−1_|*y*_*t*−1_) and the natural prior distribution (Eq 5). Depicted are two mixtures of the same components. (A) Equal mixture (solid blue line) of 50% previous posterior (dotted red line) and 50% natural prior (dotted green line). (B) Mixture (solid blue line) of 10% previous posterior (dotted red line) and 90% natural prior (dotted green line).

#### Model parameters

For the natural prior model (NPM) we used the same perceptual noise parameter (*κ*_*L*_) as in the VMF described above. Here we used an additional free parameter - the proportion of the previous posterior (*β*_1_) in the prior distribution (Eq 6). Importantly, when *β*_1_ > 0.5 (Fig 6A) the previous posterior would dominate the resulting prior and hence the model would start to approximate the VMF. Similarly, as *β_1_* approaches zero (Fig 6B) the natural prior component will dominate the prior distribution. As in our previous simulations we chose the free parameter values (*κ_Q_, β*_1_) to minimise the distance between the simulation and behavioural results observed by Fischer and Whitney (2014) (see *Model fitting and parameter optimisation* in *Supporting Information* for details).

Since the predictive prior distribution resulting from the mixture components is non-parametric we used a discrete circular filter to approximate the distributions in this simulation. The discrete filter is based on a grid of weighted Dirac components equally distributed along the circle (Kurz et al., 2016; Kurz, Gilitschenski, & Hanebeck, 2013) and was implemented with *libDirectional* toolbox for Matlab (Kurz, Gilitschenski, Pfaff, & Drude, 2015). Because in the unidimensional circular state space of orientations the quality of approximation is only given by the number of components, we felt that 10,000 Dirac components can adequately approximate a distribution of a circular variable. For details on the implementation of the filter algorithms see *Discrete circular filter with Dirac components* in *Supporting Information*.

### Mixture model

In order to model a time-decaying recency bias over several past states we modify the basic VMF so that the state transition model (*a*(·), Eq 1) predicts the next state based on a recency-weighted mixture of *m* past states:

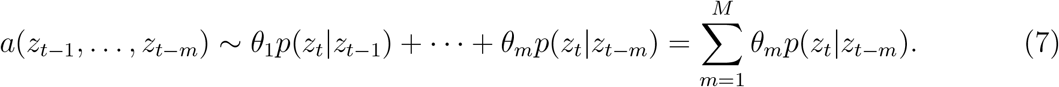

Here *θ_m_* is a mixing coefficient for the *m*-th past state. We can control the individual contribution of a past state *z_m_* to the resulting mixture distribution by defining how the mixing coefficient *θ* decays over the past states:

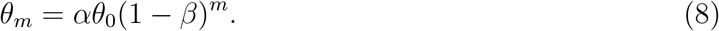

Here *β* is the rate of decrease of the mixing coefficient over the past *m* states and *α* is a normalising constant. As a result we have a decaying time window into the past *m* states defined by the rate parameter *β*. The role of the *β* parameter is to control the decrease of the mixing coefficient *θ* over past states. Fig 7A illustrates the relationship between the *β* and *θ* parameters: the bigger the *β* the faster the contribution of past states decreases and greater the proportion of most recent states to the mixture distribution (Eq 7). As *β* approaches 1, the mixture begins to resemble *z*_*t*-1_ and approximate the first-order VMF described above:

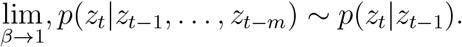

Conversely, as *β* approaches zero, all past states contribute equally to the mixture. Intuitively, *β* could be interpreted as the bias towards more recent states. Fig 7B illustrates the evolution of the latent state distribution *p*(*z*) when *β* = 0.5 and the mixing coefficient decays over the previous states.

Importantly, the mixture distribution is computed by a fixed sampling step (for details of the mixture sampling algorithm see Kalm, 2017, and *Mixture model* in *Supporting Information)*. Hence the mixture model is computationally first-order Markovian and has the same number of parameters and model complexity as the NPM described above.

In sum, we have a circular Bayesian filter where the state transition model *a*(·) is a mixture function over past *m* states (Eq 7). The proportion of the individual past states in the mixture - and therefore the effective extent of the window into the past - is controlled by the *β* parameter. As in previous models, the measurement model is identity, and both state and measurement noise (*κ_Q_* and *κ_L_*) are additive Von Mises.

**Figure 7:**
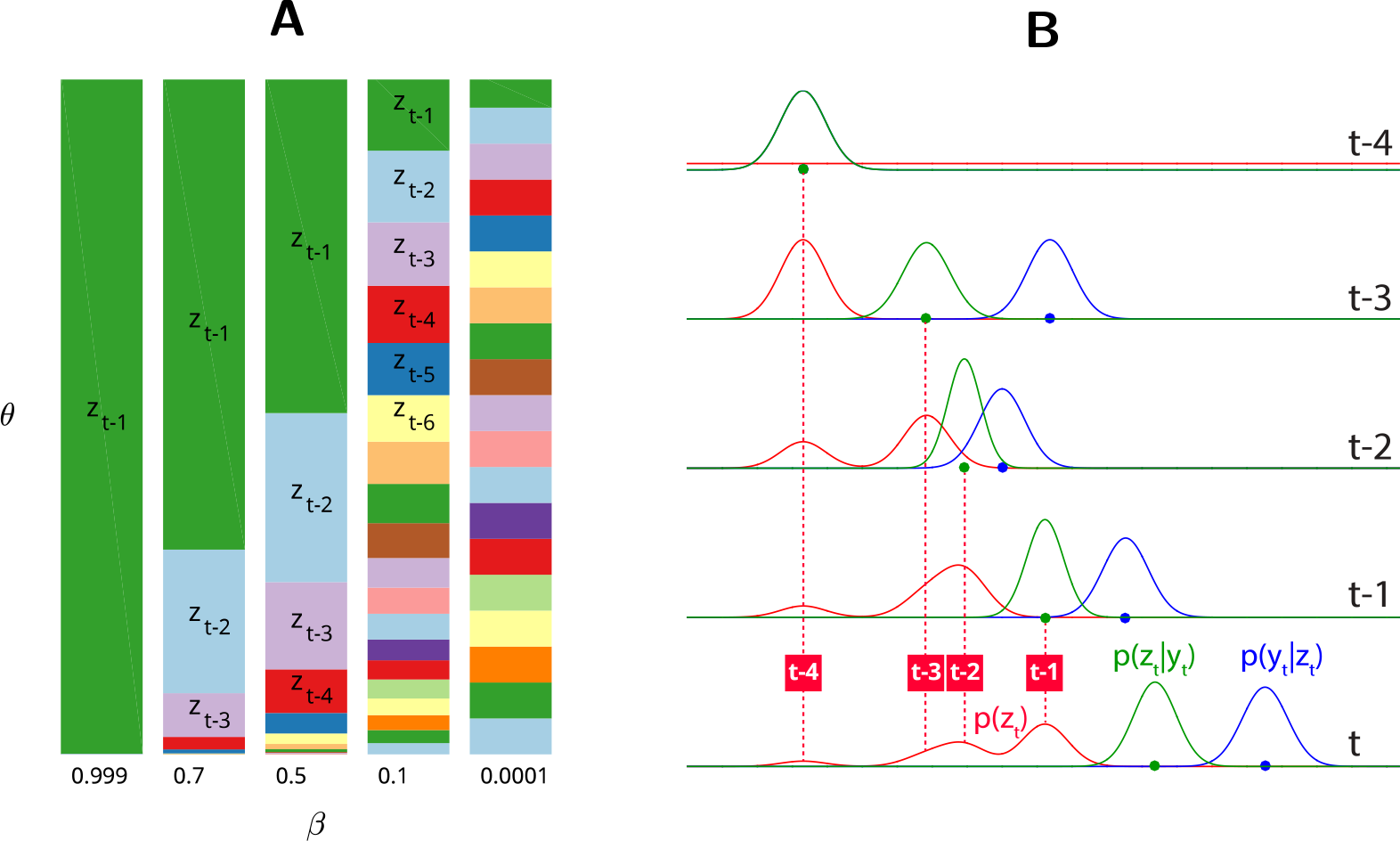
Mixture model. (A) Values of the mixing coefficient *θ* over past states (*z*_*t*-1_,…, *z_t-m_*) based on different *β* values. *θ* represents the proportion of a past state *z_t_* in the mixture distribution (Eq 7). (B) Evolution of the latent state *p*(*z*) over 4 trials.

#### Model parameters

We used the same perceptual noise parameter (*κ_L_*) as in the VMF and NPM simulations described above. The free parameters in the mixture model (MM) were the mixing coefficient hyper-parameter *β* (Eq 8) and state noise (*κ_Q_*). As with previous simulations, the free parameters were chosen to minimise the distance between the simulated data and the average observed subjects’ response (see *Model fitting and parameter optimisation* in *Supporting Information)*.

### Statistical effects of interest

In each trial, we simulated the participant’s response *k_t_* by taking a random sample from the posterior distribution: *k_t_* ~ *p*(*z_t_*|*y_t_*). We then calculated three statistical effects as follows:

(1) Distribution of errors - we fit a Von Mises distribution to the simulated errors yielding mean (*μ*) and concentration values (*κ_E_*). We then calculated the similarity between our simulated error distribution and participants average distribution by assessing the probability of simulated *μ* and κ_E_ given the distribution of participants bootstrapped 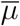 and 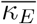.
(2) DoG recency bias curve - we fit the simulated errors with a first derivative of a Gaussian curve (DoG, see *Recency bias amplitude as measured by fitting the derivative of Gaussian* in *Supporting Information* for details), and as above, calculated the probability of the curve parameters arising from the distribution of participants’ bootstrapped parameter distributions.
(3) DoG recency bias over past three trials - we calculated the amplitude of the DoG curve peak for stimuli presented one, two and three trials back. We sought to replicate a positive but decaying recency bias over three previous stimuli.

## Results

We used all three models to simulate participants’ responses using the stimuli and experimental structure provided by the authors (824 trials with fully randomised stimuli). We sought to replicate three statistical effects observed in the behavioural experiments: zero-mean distribution of the errors, DoG-like fit of the recency bias (Fig 1B), and significant recency bias over multiple past trials (Fig 1C). See *Statistical effects of interest* in *Methods* for details.

### Von Mises Filter (VMF)

The best fitting VMF could not successfully replicate any of the three statistical effects. The distribution of errors was centred around zero but its concentration was significantly different from human data (Fig 8A-VMF, *p* = 0.026). Similarly, the maximum of the simulated recency bias was significantly removed from the human data (ca 20 deg for humans; ca 45 deg for the VMF; Fig 8B-VMF). Furthermore, it can be shown that the VMF cannot even theoretically have a maximum of the bias at less than π/4 radians (or 45 deg), which means it is incapable of replicating the DoG-like curve of the human recency bias (see *Von Mises filter properties* in *Supporting Information).* The VMF also could not replicate recency biases for stimuli presented 2 or 3 trials ago (Fig 8C-VMF). In sum, the VMF can simulate a recency bias but it is qualitatively different from human bias and only extends one trial back.

### Natural prior model (NPM)

The NPM could only partially replicate the behavioural effects. The error distribution was centred around zero but significantly different from participants’ data (Fig 8A-NPM, *p* < 0.001). However, because NPM’s prior and posterior distributions are not Von Mises it was able to capture the DoG shaped curve of the recency bias (Fig 8B-NPM). The NPM was still not able to capture either the amplitude of the recency bias or extend it back more than one previous trial (Fig 8C-NPM). This was to be expected since the NPM, like the VMF, also predicts the next state based only on the previous one (first order Markovian). In sum, the NPM was able to replicate the DoG shaped recency bias curve but only for stimuli one trial back. Furthermore, the error distributions simulated by the NPM were significantly different from participants’ average with variance of the response reduced by approximately twofold.

**Figure 8:**
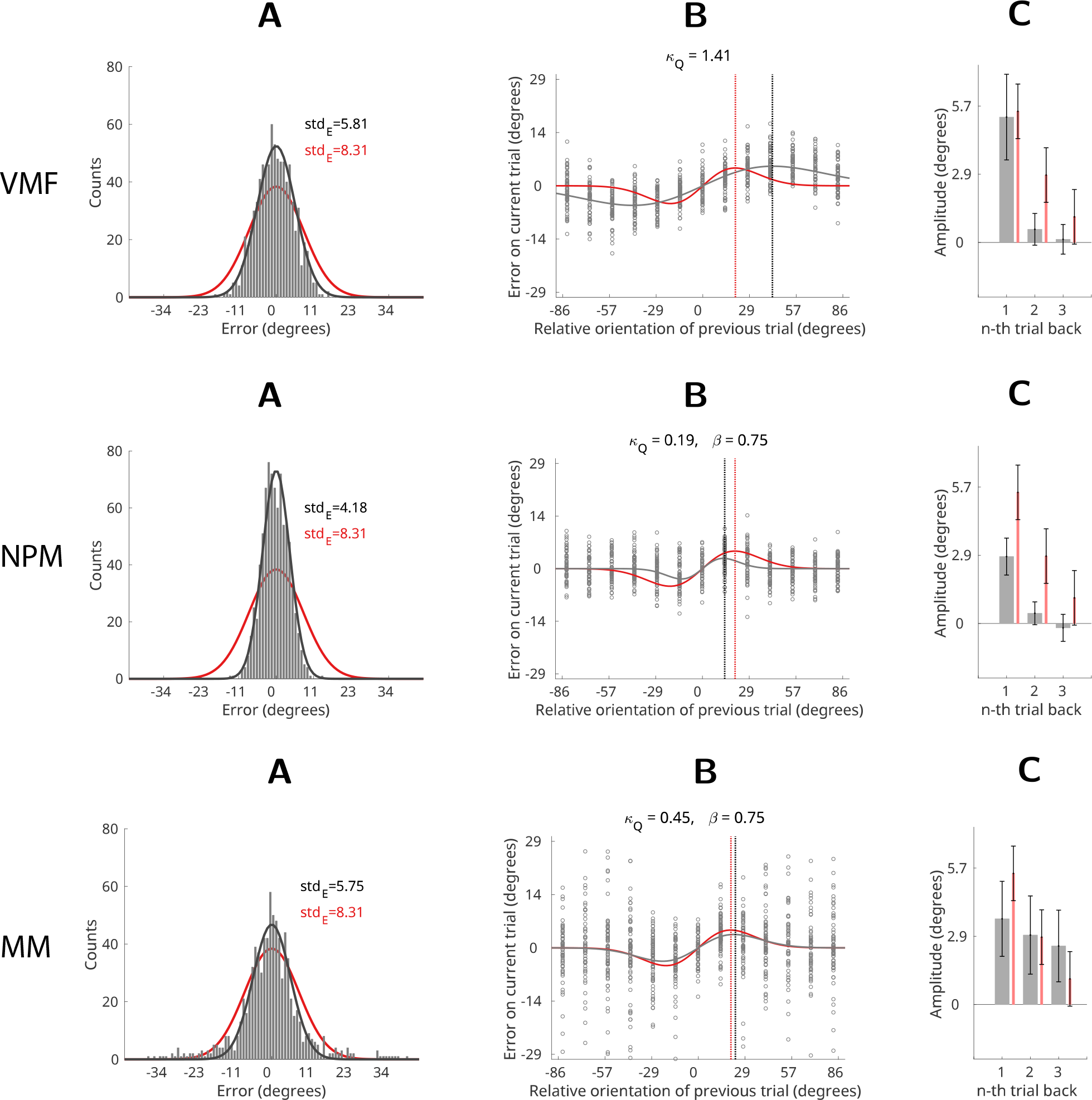
Results. Comparison of model simulation results (black) with human data from Experiment 1 (red, Fischer & Whitney, 2014). (A) Error histograms: black - model simulation, red - average human participant. Solid lines depict Von Mises fits to error distributions. (B) Recency bias: black circles show errors of the simulated responses. Solid black line shows a DoG curve fit to the simulated errors, red line shows the human recency bias (average DoG fit to human errors). Dotted vertical lines show the location of the maxima of the recency biases (black - model; red - human participants). (C) Average recency bias amplitude computed for stimuli presented one, two and three trials back from the present. Grey bars - model; red bars - human participants. Error bars represent ±1 standard deviation of the bootstrapped distribution.

#### Mixture model

The mixture model was able to successfully simulate all three statistical effects of interest: the distribution of errors was not significantly different from the participants’ data (Fig 8A-MM, *p* = 0.23); the recency bias fit the DoG-shaped curve (Fig 8B-MM), and a significant recency bias was evident over multiple past states (Fig 8C-MM). Importantly, the best-fitting *β* parameter value for the mixture model was *β* = 0.75, which effectively sets the time window for the mixture distribution at 3-4 past states (see Fig 7A, column 2, *β* = 0.7).

## Discussion

In this paper we investigated the internal model of the environment which leads people to show a recency bias.

First, we showed that participants cannot be using a simple Bayesian filter that predicts the orientation on the current trial based only on the previous one. Furthermore, we showed that a first-order identity model is theoretically incapable of producing the recency bias observed in the orientation estimation task. This suggests that previous proposals that a simple first-order Bayesian model (such as Kalman or Von Mises filters) could explain the temporal continuity over trials and hence the recency bias (Burr & Cicchini, 2014; Wolpert & Ghahramani, 2000; Rao, 1999) are misplaced. Second, we showed that a more complex model, where participants use the natural statistics of the environment in addition to the previous stimulus, is similarly incapable of simulating the recency bias. Although such an approach is significantly better at replicating the DoG like shape of the recency bias curve it still lacks a mechanism to extend the bias beyond the most recent state. Finally, we showed that a model where the prediction about the next stimulus incorporates information from multiple past orientations can successfully simulate all aspects of the recency bias. Specifically, the participant’s model of the environment is assumed to be a mixture of multiple past states so that more recent states contribute more than older ones.

The classical Bayesian interpretation of our results suggests that the recency bias is a result of model mismatch: people infer an incorrect model for data resulting in suboptimal inference. This view posits that people are either incapable of recognising randomness or inevitably assume a model for the data since it is an efficient strategy for the natural environment, where random data is rare (Bar-Hillel & Wagenaar, 1991). Specifically, if a recency-weighted prediction works well in the natural environment, where temporal continuity prevails, people would also wrongly apply that model to random data. However, a more parsimonious explanation exists. Perhaps, rather than inferring the wrong model (out of many models that might be inferred), the recency bias may simply be a consequence of the way past experiences are represented in memory.

This can be made explicit in the framework of Bayesian filtering: the prediction for the next state is calculated by applying a state transition function *a*(·) to m past states:

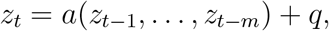

where *q* is state noise. According to the model mismatch explanation the state transition function *a*(·) is the recency-weighed mixture function (Eq 7). Importantly, this assumes that all data from past *m* states is potentially available for the state transition function *a*(·) to generate a prediction. Participants suboptimal behaviour is hence caused by applying the mixture model to data representing past *m* states. However, exactly the same prediction would be generated if the state transition function *a*(·) would not perform any transform at all - is identity - but the data from the past *m* states is itself a recency weighed mixture. This is a more realistic interpretation since the former hypothesis assumes unlimited storage for past experiences. Similary, the latter interpretation does not require any model selection at all (out of possibly infinite models) and is hence a more parimonious view.

Consider what happens when the model of the environment is unknown and needs to be inferred in real time: for random data, such model inference is always bound to end in failure as no model can explain, compress, or more efficiently represent random input. The most efficient representation of a random latent variable is the data itself and not data plus model. In other words, people might not be applying the wrong model to the data, rather they may be failing to apply any model at all. The recency-weighted bias over multiple past states instead reflects the observer’s representation of the past. This is in agreement with previous proposals that stimulus representation in visual estimation tasks might include partially ‘overwriting’ previous representations with newer ones (Matthey, Bays, & Dayan, 2015). Note that abandoning Bayesian inference altogether by simply ignoring the previous states would actually result in optimal performance in the task with random data. However, this strategy would immediately run into trouble should a pattern begin to emerge in data which is initially random.

Therefore we propose that instead of having the data and just applying a ‘wrong’ model to it - a classic case of Bayesian model mismatch - the recency bias emerges because participants are continuously and unsuccessfully attempting to infer a model based on previous states. In formal terms, the state transition model contains no information (it is identity) and hence the predictive prior distribution simply reflects the observer’s representation of the past.

In sum, our results indicate that the recency bias that appears when participants are confronted with random data must be driven by a mixture of past states. We suggest that the most parsimonious explanation of our results is that participants fail to infer a model for the data and fall back on treating the internal representation of the data itself as the best prediction for the future.

## Supporting Information

### Bayesian orientation estimation

We can model participants behaviour on the orientation estimation task (Fig 1) as probabilistic inference by combining their prior expectations about the orientation of the Gabors with immediate sensory evidence on a given trial (see any of Fiser, Berkes, Orbán, & Lengyel, 2010; Ghahramani, 2015; Pouget, Beck, Ma, & Latham, 2013; Griffiths, Chater, Kemp, Perfors, & Tenenbaum, 2010 for a primer on human probabilistic infererence).

Hence, we can model the participants internal representation (*z*_*t*_) given the orientation of the presented Gabor (*y*_*t*_) on trial *t* as Bayesian inference:

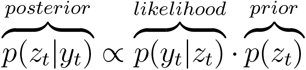

The posterior distribution *p*(*z*_*t*_|*y*_*t*_) represents a participant’s estimate of the orientation on trial *t* and their response can be thought of as a sample from the posterior distribution. Since orientation is a circular variable we need to use a probability distribution wrapped on a circle to represent variables of interest (here we use a Von Mises distribution, Fig 2A). We model the sensory evidence, or the likelihood distribution, as Von Mises noise around the value of the stimulus *y_t_* on trial:

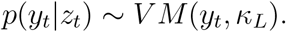

We fix the value of the likelihood noise parameter *κ*_*L*_ for every trial and derive it from the participants’ average just noticeable difference (JND) as derived with Experiment 3 in Fischer and Whitney (2014) (see *Measurement noise concentration parameter* in *Supporting Information* for details).

#### Bayesian filtering

We assume that at every time-step *t* observation *y_t_* corresponds to a latent variable *z*_*t*_ (internal representation of orientation), which over time forms a Markov chain, giving rise to the latent state space model. Hence, the state of the latent variable *z_t_* is inferred at every time step t using the Bayes theorem based on the previous state *z*_*t*−1_ and current observation (sensory information) *y_t_*:

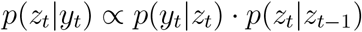

Assuming that noise in the internal representation and observation is additive, then the evolution of the latent variable *z* is predicted by the state transition model and the likelihood of the observations *y* are given by the measurement model:

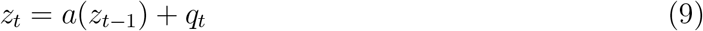

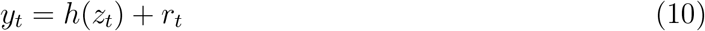

Assuming that the state transition and measurement models (*a* and *h*) are arbitrary but known functions and *q_t_* and *r_t_* are additive noise, we can carry out Bayesian inference at every time step by first predicting the next state of the latent variable given the past state (Eq 9) and then updating this prediction when observable data is measured (Eq 10) to give us a Bayesian posterior distribution of the latent variable (see Sarkka, 2013; Bishop, 2006, for a detailed description of state space models).

This two-step process is called *Bayesian filtering*. If the evolution of the state-space is linear and the noise is Gaussian then the optimal probabilistic state space model is the Kalman filter (Kalman & Bucy, 1961). Kalman filters have been extensively used for problems where the future state of the environment can be predicted from just the previous few states with sufficient accuracy, such as in object tracking or phoneme recognition. An in-depth mathematical explanation of the Kalman Filter, and how it is derived from the Bayesian estimation problem can be found in any good book covering digital signal processing, for example in Sarkka (2013). However, Gaussian distributions are not appropriate when the latent state space is wrapped on a circle. A circular analogue of the Kalman filter is the Von Mises filter.

### Measurement noise concentration parameter

Participants’ psychometric functions were estimated in Fischer and Whitney (2014) as just noticeable differences (JND) by using a two-interval forced-choice task (Experiment 3 in Fischer & Whitney, 2014). We converted the JND values (*σ_P_*) to the measurement concentration parameter *κ_L_* as JND relates to the standard deviation (*σ*) of the normal distribution:

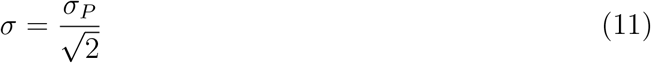

and since the concentration of the Von Mises distribution can be approximated as κ = 1/*σ*^2^ we get

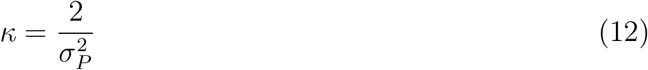

The mean JND (*σ_P_*) across subjects was 5.39° (Fischer & Whitney, 2014), so we used *σ* = 3.8113° and *κ_L_* = 0.0688.

### Product of two von Mises distributions

Given two von Mises probability density distributions *p*(*x*; *μ*_1_, *κ*_1_) and *p*(*x*; *μ*_2_, *κ*_2_) the resulting product is an unnormalised von Mises distribution (Murray & Morgenstern, 2010):

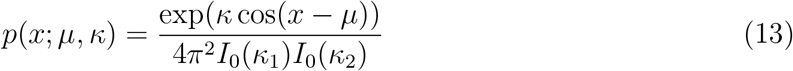

where the mean *μ* and concentration *κ* are respectively:

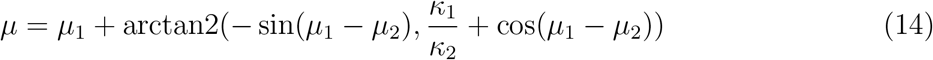

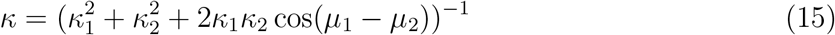

### Von Mises filter properties

In the Von Mises filter (VMF) the recency bias at any trial *t* is given as the distance which posterior mean 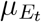 has moved away from the presented stimulus *y_t_* towards some previous stimulus value *y_t-n_*. Since the posterior in VMF is a product of two individual VM distributions (likelihood and prior) such distance (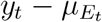, recency bias on trial *t*) can be computed analytically by evaluating 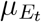 as Eq 14:

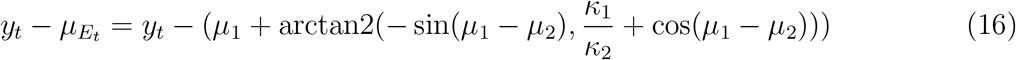

Since *y_t_* is the mean of the likelihood and hence *μ*_1_ of the first Von Mises components, we can fix the prior mean (*μ*_2_ of the second Von Mises component) to zero, and hence assign:

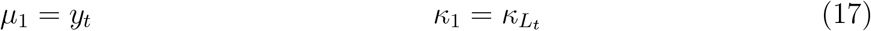

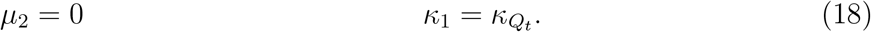

Making these replacements in Eq 16 gives us:

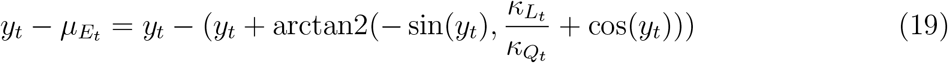

which after some rearranging becomes:

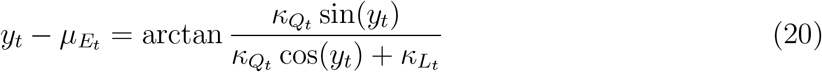

It follows that the estimation error (Eq 4) is only dependent on two parameters: (1) distance between the prior mean and stimulus (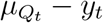, x-axis on Fig 5); and (2) ratio of likelihood to prior concentration (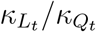, differently coloured lines on Fig 5B). Such mapping of all possible shapes of the recency bias reveals several interesting findings.

First, in order to observe a recency bias the concentration of the likelihood has to be greater than the concentration of the prior: 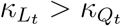. Conversely, when 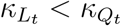 the posterior mean will be closer to the prior mean than to the stimulus and the bias curves on Fig 5 will be inverted. In other words, likelihood concentration has to be on the average greater than prior for a participant’s response error distribution to be centred around zero. In general terms this means that perceptual noise in the task has to be smaller than uncertainty about the next stimulus.

Second, one would intuitively expect the estimation error (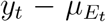, y-axis), and hence the recency bias, to be greatest when the distance between the prior and stimulus 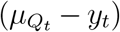 is greatest (x-axis maxima and minima on Fig 5B). However, as shown in Fig 5B for various values of 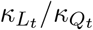, this is not the case because of the circular nature of Von Mises. As the distance between the prior mean and stimulus approaches maximum (antipodean angle) the influence of the prior decreases so that the mean of the posterior tends back towards the stimulus. At maximum distance (±π/2 on the x-axis on Fig 5B) the influence of the prior is zero and the posterior mean is equal to the stimulus.

Finally, and most importantly, no value of 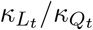 can even theoretically yield a DoG-like recency bias shape as observed in Fischer and Whitney (2014) (Fig 1B): the Von Mises recency bias cannot have maxima or minima between −π/4 and π/4 (see *Minima and maxima of the recency bias in Von Mises filter* below for details). In other words, when Von Mises distributions are used for Bayesian inference the recency bias always peaks more than halfway through the x-axis. Contrastingly, DoG curves fitted to participant data in Fischer and Whitney (2014) peak close to zero and between ±π/4 (Fig 1B). In sum, a DoG-shaped recency bias is not even theoretically possible if participants use Bayesian inference and Von Mises distributions for orientation estimation.

### Minima and maxima of the recency bias in Von Mises filter

We find the extrema of Eq 4 by setting its first derivative to zero:

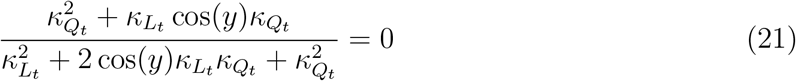

which gives us maximum and minimum at

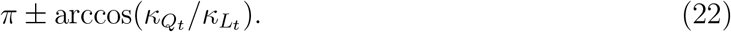

In order to observe a positive recency bias the likelihood concentration has to be greater than prior’s 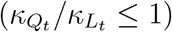 and therefore arccos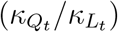 can only take values between [0,π/2].

Finally, we need to convert this result to the stimulus space used in the behavioural experiments - the Von Mises distribution is wrapped on a full circle [0,2π], whilst in the orientation estimation task stimulus values were wrapped on the top half of the circle [−π/2,π/2] (Fischer & Whitney, 2014). Eq 22 wrapped between ±π/2 gives ±π/4 as the new limits to the extrema.

### Recency bias amplitude as measured by fitting the derivative of Gaussian

The errors as a function of between-trial orientation distance (Fig 1B) were fitted with a first derivative of a Gaussian curve (DoG) as given by Fischer and Whitney (2014):

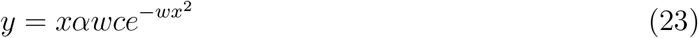

where *x* is the relative orientation of the previous trial, *α* is the amplitude of the curve peaks, *w* scales the curve width, and *c* is the constant 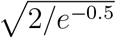 which rescales the curve so that the *α* parameter numerically matches the height of the positive peak of the curve for ease of interpretation: the amplitude of serial dependence (*α*) is the number of radians that perceived orientation was pulled in the direction of the previous stimulus.

### Model fitting and parameter optimisation

To evaluate how well the simulated recency bias fit with the observed behavioural data we measured the distance between the simulated recency bias DoG curve to the one observed in the behavioural experiments (Fischer & Whitney, 2014). The distance between the simulated and observed curves was measured as the sum of squared errors (SSE) across orientations bins. Fig 0.1 displays the average DoG curve recency bias fit across the participants. To find the value of free parameters (latent state noise and mixing coefficient) that yielded recency biases most similar to behavioural results we performed a grid search across parameter space. We used state noise *κ_Q_* values from 48 to 480 (step size *log*(*x*)) and *β* values from 0.2 to 0.9 (step size 0.05). We used the parameter values which yielded the lowest SSE to the average participants DoG curve (Fig 0.1).

### Natural prior

Here we used the average participants’ prior as reported in (Girshick et al., 2011) as the natural prior distribution. This was modelled as a mixture of two von Mises probability density distributions with means fixed at 0 and π radians respectively.

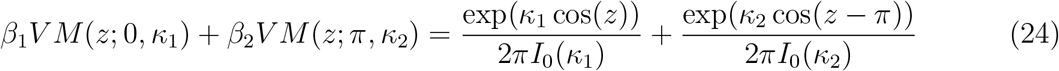

**Figure 1:**
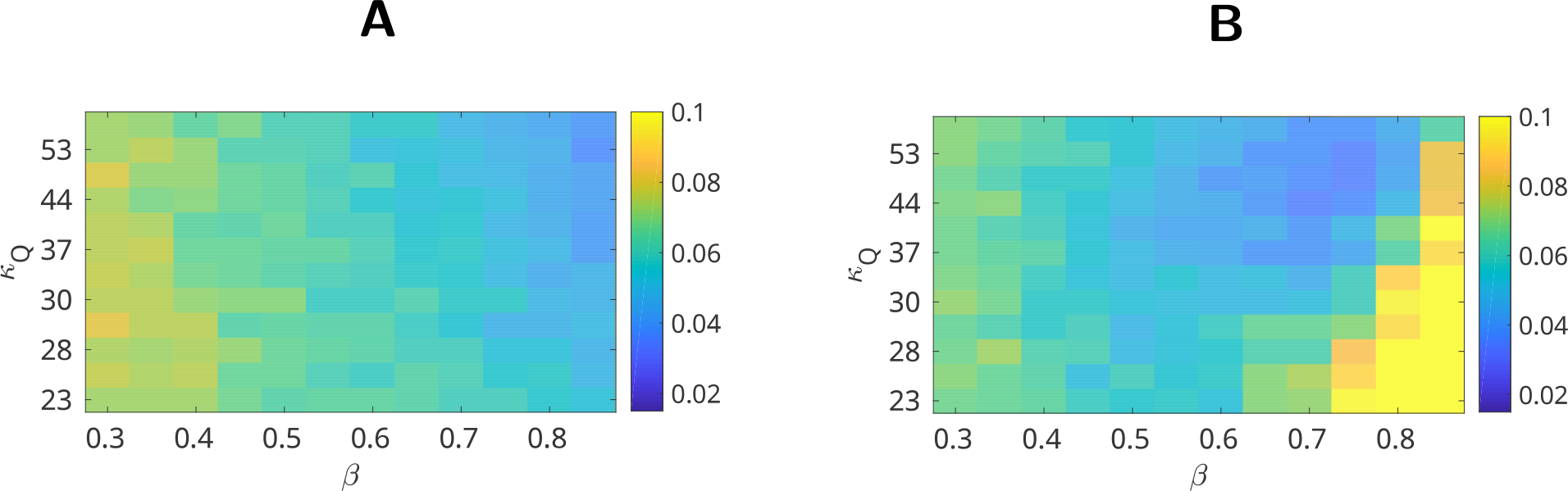
Parameter search grid results. Sum of squared errors (SSE) between simulated recency bias curves and the observed human bias. SSE is displayed as a function of state noise concentration parameter (*κ_Q_*, y-axis) and mixture coefficient (*β*, x-axis). (A) Natural prior mixture. (B) Von Mises mixture.

We used Matlab’s nonlinear and derivative-free model fitting function *fminsearch* with 10^8^ iterations and an ordinary least squares cost function to estimate the values of mixing coefficients (*β*_1_ = 0.0215, *β*_2_ = 0.0178) and von Mises concentrations (*κ*_1_ = 0.8365 *κ*_2_ = 0.8728). The resulting mixture and its fit with data from (Girshick et al., 2011) is depicted on Fig 3C.

### Discrete circular filter with Dirac components

A detailed derivation of the circular filter using Dirac mixtures can be found in Kurz et al. (2013, 2016). Briefly, a wrapped Dirac mixture with *L* components and Dirac positions *β*_1_,…, *β_L_* ∊ [0, 2*π*] is defined as:

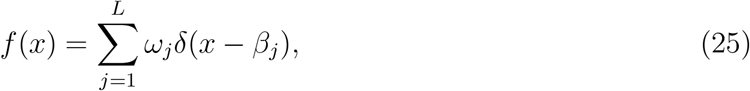

where *ω_j_* are weighting coefficients and 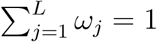. A Von Mises distribution can be approximated by a wrapped Dirac mixture

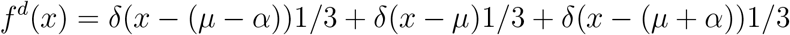

by calculating *μ* as the circular mean

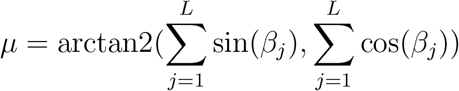

i. e., the argument of the first circular moment, and by matching the first circular moment to obtain *κ*.

### Mixture model

A detailed derivation of the mixture model and the sampling algorithm can be found in Kalm (2017); Raftery (1985). What follows is a brief overview of the approach.

The latent state *z* can be modelled as the mixture of it’s past states by using a mixture state transition function (see Berchtold & Raftery, 2002; Raftery, 1985, for details). This method considers the effect of the each of the past m states separately. Specifically, the conditional probability distribution *p*(*z_t_*|*z*_*t*−1_,…, *z_t-m_*) is modelled by a mixture distribution of past *m* states:

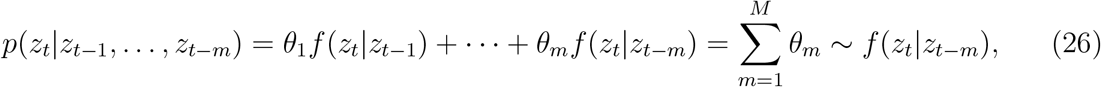

where *θ* is a mixing coefficient so that

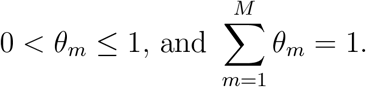

We assume that the mixing coefficient *θ*, declines over *m* time steps as given by some decay function *ϕ* and the rate of decay parameter *β*:

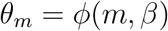

Here we use an exponential decay function *ϕ*, so that:

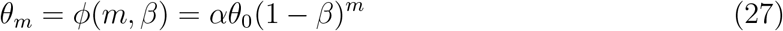

where *β* is the rate of decrease (0 ≤ *β* < 1) and *α* normalising constant. Substituting *θ_m_* into Eq 26 gives:

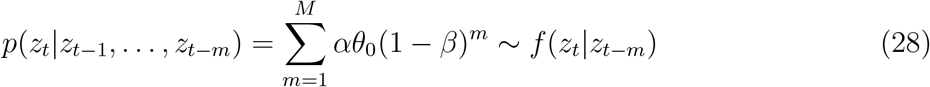

In order to limit the computational cost of performing inference at every time step we represent the distribution of latent variable *z* with a fixed number of samples *L*. As a re-sult the proportion of samples assigned to a particular component of the mixture distribution (representing a past state) is determined by the mixing coefficient *θ*:

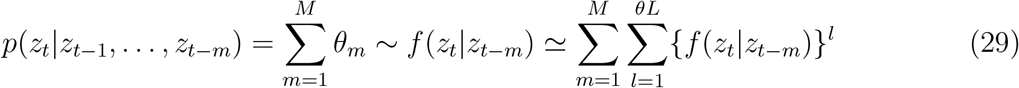

where *L* is the total number of samples, *θL* is rounded to the nearest integer, and {*f* (*z_t_|z_t-m_*)}^l^ is a set of *l* samples drawn from *p*(*z_t_|z_t-m_*}.

The property of constant number of samples *l* for every *m*-th component of the mixture at any time-step ({*z_t_|z_t-m_*}^l^} is important since it greatly simplifies the approximation of the mixture distribution (Eq 28) algorithmically. If at every time-step *t* we take a fixed proportion *β* from the existing mixture distribution and reassign those samples to represent the most recent component, then after *m* steps we end up with the same proportion of components as given by (Eq 28). It follows that a mixture distribution of past *m* states with exponentially decaying proportions of past *m* components can be approximated by sampling from just *f* (*z_t_*|*z*_*t*−1_) and the previous state of the mixture {*z*_*t*−1_} at every time step *t*.

## Acknowledgments

We would like to thank Jason Fischer for sharing with us the experimental data from Fischer and Whitney (2014).

